# Repurposing screen highlights broad-spectrum coronavirus antivirals and their host targets

**DOI:** 10.1101/2021.07.14.452343

**Authors:** Sibylle Haid, Alina Matthaei, Melina Winkler, Svenja M. Sake, Antonia P. Gunesch, Jessica Rückert, Gabrielle Vieyres, David Kühl, Tu-Trinh Nguyen, Lisa Lasswitz, Francisco Zapatero, Graham Brogden, Gisa Gerold, Bettina Wiegmann, Ursula Bilitewski, Mark Brönstrup, Thomas F. Schulz, Thomas Pietschmann

**Author notes:** Corresponding author: Thomas Pietschmann, Institute for Experimental Virology, Twincore -Centre for Experimental and Clinical Infection Research; Feodor-Lynen-Straße 7, 30625 Hannover, Germany. These authors contributed equally to this work (Sibylle Haid, Alina Matthaei, Melina Winkler).

## Abstract

Libraries composed of licensed drugs represent a vast repertoire of molecules modulating physiologic processes in humans, thus providing unique opportunities for discovery of host targeting antivirals. We interrogated the ReFRAME repurposing library with 12,993 molecules for broad-spectrum coronavirus antivirals and discovered 134 compounds inhibiting an alphacoronavirus, mapping to 59 molecular target categories. Dominant targets included the 5-hydroxytryptamine receptor and dopamine receptor and cyclin-dependent kinase inhibitors. Counter-screening with SARS-CoV-2 and validation in primary cells identified Phortress, an aryl hydrocarbon receptor (AHR) ligand, Bardoxolone and Omaveloxolone, two nuclear factor, erythroid 2 like 2 (NFE2L2) activators as inhibitors of both alpha- and betacoronaviruses. The landscape of coronavirus targeting molecules provides important information for the development of broad-spectrum antivirals reinforcing pandemic preparedness.

## 1. Introduction

Three genetically diverse coronaviruses have caused two epidemics and one pandemic in the past two decades. These emerging pathogens have transmitted zoonotically from animal hosts. They exhibit differential transmission characteristics and variable degrees of pathogenicity in humans. Within the subfamily of *Coronavirinae*, harboring five genera, these viruses map to two distinct subgenera of the genus *Betacoronavirus*: the subgenus *Sarbecovirus* (SARS-CoV and SARS-CoV-2) and the subgenus *Merbecovirus* (MERS) (Coronaviridae Study Group of the International Committee on Taxonomy of, 2020) (Fig.1A). Additional coronaviruses have been circulating among humans and usually cause mild respiratory and intestinal disease. These include HCoV HKU1 and HCoV OC43 of the subgenus *Embecovirus* within the genus *Betacoronavirus* and the alphacoronaviruses HCoV NL63 (subgenus *Setracovirus*) and HCoV 229E (subgenus *Duvinacovirus*). Related coronaviruses from these branches of phylogeny circulate in various mammals, including bats, rats, mice, and pigs, which live in close proximity to humans. It is unclear which viral determinants control zoonotic transmission to humans. Moreover, we lack information about which animals host coronaviruses with the potential to spill over to and spread among humans. Therefore, we are facing unpredictable risks of entering additional pandemics in the future initiated by new and genetically diverse coronaviruses. Multiple vaccine candidates for prevention of SARS-CoV-2 infections are under clinical development, the first ones have been approved and large-scale vaccination campaigns have recently started. However, there is uncertainty if any of these vaccine candidates would prevent infections by other coronaviruses. Further to this, natural immunity to circulating CoVs and SARS-CoV is usually relatively short-lived (Callow et al., 1990; Cao et al., 2007; Wu et al., 2007). Bamlanivumab (LY-CoV555), a viral spike protein-targeting human neutralizing antibody (Chen et al., 2020), received FDA emergency use authorization for treatment of recently diagnosed mild to moderate COVID-19 disease in high-risk patients. However, whether this or other antibodies under clinical development would be efficacious for treatment of other emerging coronaviruses is unknown (DeFrancesco, 2020). Remdesivir, a broad-spectrum nucleoside analogue, inhibits replication of diverse coronaviruses and was licensed for treatment of severe COVID-19. However, its efficacy for treatment of SARS-CoV-2 infections appears to be moderate on the basis of differential outcomes of clinical trials and the WHO no longer recommends its use in COVID-19 patients (Young et al., 2020). Therefore, novel therapeutic options for diverse coronaviruses are needed.

Viruses are intracellular parasites and take advantage of cellular co-factors in multiple ways. Some of these host factors are shared between diverse coronaviruses (Schneider et al., 2021). Interference with these shared host factor dependencies is an attractive route for development of broad-spectrum antiviral therapies. However, comprehensive screens for druggable pathways and host targets as well as drug candidates with broad anti-coronavirus activity are lacking. The ReFRAME library accumulates 12,993 small molecules (Janes et al., 2018). Around 68% of these compounds either are licensed for use in humans or are in advanced stages of clinical development (Fig. 1B). Disease annotations of these molecules are dominated by cancer (43%), central nervous system (CNS, 23%), and cardiovascular/respiratory (20%), so that 86% of molecules in the ReFRAME library target physiological processes in humans (https://reframedb.org, April 2020). This composition renders the library a unique resource for identification of molecules targeting host pathways and host factors with potential for development of broad-spectrum antiviral therapies. Screening of this library with SARS-CoV-2 revealed interesting drug repurposing candidates (Riva et al., 2020). Here, we provide a comprehensive overview of drugs and drug-like molecules with therapeutic potential against genetically diverse coronaviruses including the host factors and cellular pathways they target.

**Fig. 1.**
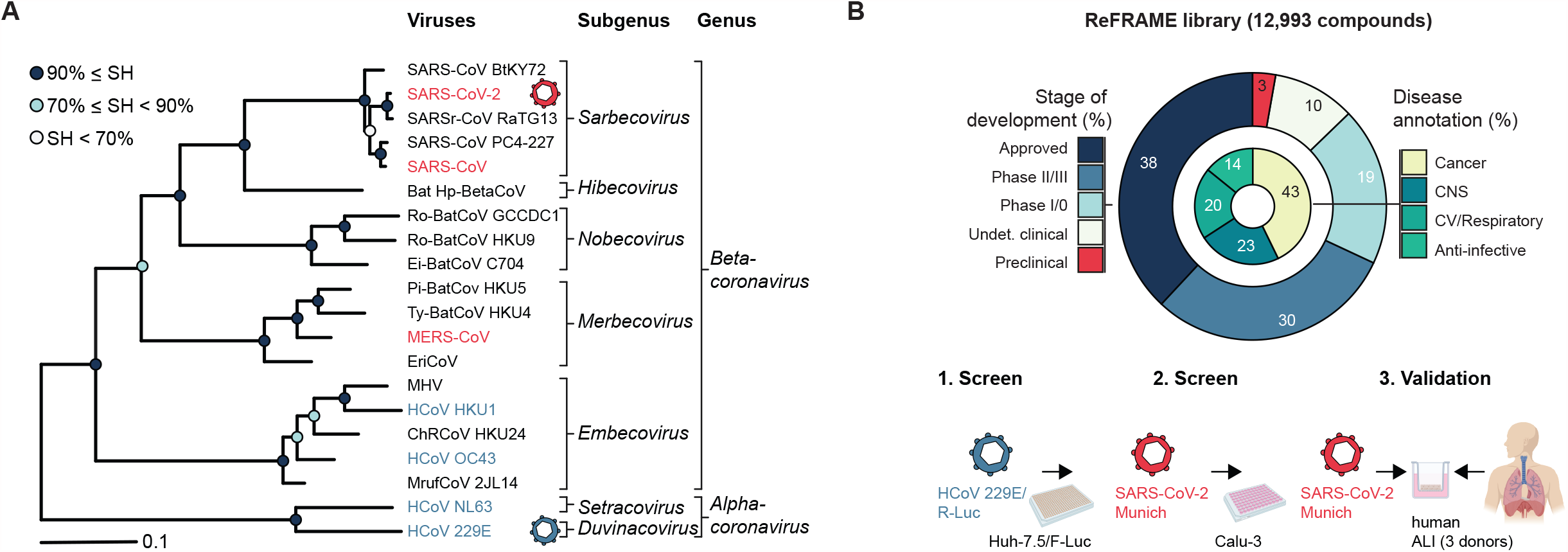
Genetic diversity of human pathogenic coronaviruses, ReFRAME library composition and screening workflow. **(A)** A section of the coronavirus phylogeny based on an IQ-TREE maximum-likelihood tree showing representative viruses of sixteen species representing all fourteen species of the genus Betacoronavirus and two of the genus Alphacoronavirus (adapted from 1). Branch support was estimated using the Shimodaira-Hasegawa (SH)-like approximate likelihood ratio test with 1,000 replicates. **(B)** Composition of the ReFRAME drug repurposing library and schematic of the screening workflow for identification and validation of broad-spectrum coronavirus antivirals (created with BioRender.com).

## 2. Material and methods

### 2.1. Cell lines

The following cell lines were used for this study: Vero cells (*Chlorocebus aethiops*, ATCC-CCL-81; Lot 58484194), Vero E6 cells (*Chlorocebus aethiops)* and Calu-3 cells (*Homo sapiens sapiens*) (both from Stefan Pöhlmann), Huh-7.5 cells (*Homo sapiens sapiens*, Charles M. Rice), MRC-5 cells (*Homo sapiens sapiens*, Volker Thiel), 16HBE140 (*Homo sapiens sapiens*, Gert Zimmer), A549 cells (*Homo sapiens sapiens*, ATCC-CCL-185; Lot 59239596), HEp-2 (*Homo sapiens sapiens*, ATCC-CCL-23; Lot 58978772), A427 (*Homo sapiens sapiens*, cell line services CLS# 300111; Lot 300111-612).

### 2.2. HCoV 229E screening

Huh-7.5/F-Luc cells were seeded in black 384-well plates with a flat clear bottom (Corning #3764) at a density of 3×10^3^ cells/well in phenol-red free Dulbecco’s modified Eagle medium (DMEM; Gibco #31053-028) supplemented with 2 mM L-glutamine (Gibco #25030024), 100 µg/mL streptomycin and 100 U/mL penicillin (Gibco; #15140122), 10% fetal calf serum (FCS, Capricorn Scientific #FBS-11A) and non-essential amino acids (NEAA; Gibco #11140035) one day prior to infection. Cells were infected with a HCoV 229E *Renilla* luciferase reporter virus with an MOI of 0.1 in presence of a final drug concentration of 5 µM using a Beckman Coulter I5 automated workstation. Blinding was done by Calibr by providing ID numbers instead of compound names. The SCRIPPS Research Institute quality controlled the compounds contained in the ReFRAME library via liquid chromatography mass spectrometry (LC-MS) and Nuclear magnetic resonance spectroscopy ^1^ H-(NMR) to acquire a purity of ≥ 95% (Janes et al., 2018). Following incubation for 48h at 33°C, luciferase activity was determined using the dual-glo luciferase assay system (Promega, #E2920) and a GloMax machine (Promega; GM3000; software version 3.1.0) according to the instructions of the manufacturer. In brief, 60 µl of cell culture fluid was removed from each well to obtain a final volume of 20 µl per well prior to addition of 20 µl of Dual-glo luciferase assay reagent, incubation for 20 min at room temperature and subsequent firefly luciferase measurement (0.5 sec detection time). Firefly luminescence was quenched by addition of 20 µl per well Dual-glo stop & glo reagent. After additional 20 min incubation time at room temperature, *Renilla* luminescence was analyzed (0.5 sec detection time).

### 2.3. Virus stock

Vero cells (ATCC-CCL-81; Lot 58484194) were cultured in T75 flasks in Advanced Minimal Essential Medium supplemented with 1% NEAA, 100 U/mL penicillin, 100 μg/mL streptomycin, 2 mM L-glutamine and 10% FCS at 37°C (5% CO2). At 50-80% confluence, Vero cells were inoculated with 500 µL of SARS-CoV-2 (strain SARS-CoV-2/München-1.2/2020/984; p3) in a total of 10 mL of Advanced MEM and incubated at 37°C (5% CO2) under BSL-3 conditions adapted for infectious respiratory viruses. As soon as cytopathic effects were visible, mostly at 96 hours post infection (hpi), supernatant was collected, centrifuged at 1,000 x g for 10 min and stored in aliquots at -80°C. Stocks were titrated on Vero and Calu-3 (Finkbeiner et al., 1993) cells (ATCC Cat. #HTB-55 and RRID:CVCL_0609, respectively), cultured in DMEM containing 1% NEAA, 100 U/mL penicillin, 100 μg/mL streptomycin, 2 mM L-glutamine, 10% FCS. At 72 hpi cells were fixed with 10% formalin for 30 min and stained with crystal violet. Viral titer (TCID50/mL) were quantified based on the Spearman and Kärber method using a calculator developed by Marco Binder, Heidelberg University. HCoV 229E/R-Luc stocks were prepared by inoculation of Huh-7.5 cells. After 48h of incubation at 33°C, virus-containing supernatant was harvested and cleared from cell debris by centrifugation at 1,000 x g for 10 min prior to storage at -80°C.

### 2.4. SARS-CoV-2 infection in Calu-3 cells and primary human airway epithelial cells

For infection with SARS-CoV-2, Calu-3 cells (Finkbeiner et al., 1993) were seeded in 96-well plates 48h prior to virus inoculation at a density of 5×10^4^ in DMEM (Gibco #41965039) supplemented as described above. Two hours prior to infection, cells were treated with respective compounds in a volume of 150 µl per well at 37°C. Infection with SARS-CoV-2 was conducted with an MOI of 68.2 TCID50/well for Calu-3 cells, corresponding to 1.36×10^-3^ TCID50/cell and resulting in a final volume of 200 µl/well. For infection of differentiated primary human airway epithelial cells cultured under air liquid interface conditions cells were inoculated with 1.16×10^5^ TCID50/well or 1.58×10^3^ TCID50/well of SARS-CoV-2 for 1h at 37°C in presence of 1 or 5 µM of the respective compound. As control, heat-inactivated inoculum (70°C, 15 min) was used. Following incubation for 48h at 37°C, supernatant was harvested from Calu-3 cells and inactivated by addition of lysis buffer (Maxwell 16 viral total nucleic acid purification kit, Promega #AS1150) complemented with proteinase K. For analysis of SARS-CoV-2 production of primary human lung epithelial cells, cells were washed twice with a total of 200 µl of Hank’s Balanced Salt Solution (HBSS, Gibco; #14175129) and lysed as described above. Primary human airway epithelial cells were homogenized in homogenization solution supplemented with 1-thioglycerol and lysed by addition of lysis buffer (Maxwell 16 LEV simplyRNA cells kit; Promega #AS 1270). Prior to RNA extraction, all samples were heat-inactivated at 70°C for 15 min. For fluorescence imaging Calu-3 cells were fixed in 10% formalin for 30 min.

### 2.5. RNA extraction and qRT-PCR analysis

RNA from lysed and heat-inactivated samples was purified according to the instructions of the manufacturer using Maxwell 16 viral total nucleic acid purification kit (Promega, #AS1150) or Maxwell 16 LEV simplyRNA cell kit (Promega; #AS1270), respectively. Purified RNA was eluted in 50 µl nuclease-free water and stored at -80°C. Quantitative RT-PCR was performed using a LightCycler 480 machine (Roche, software version 1.5.1.62), primers and probe specific for SARS-CoV-2 RdRP (TibMolBiol; #53-0777-96) and a LightCycler Multiplex RNA Virus master kit (Roche; #07083173001) according to the instructions of the manufacturer. Quantification of copy numbers was done using an in-run standard curve. For GAPDH detection, the following primer and probes were used (S-GAPDH, 5′-GAA GGT GAA GGT CGG AGT C -3′; A-GAPDH, 5′-GAA GAT GGT GAT GGG ATT TC -3′; probe: LC640-CAA gCT TCC CgT TCT CAg CCT –BBQ;

TibMolBiol). Means of double measurements for each sample are given.

### 2.6. Fluorescence microscopy and quantitative image analysis

Following infection, cells used for immunofluorescence staining were fixed with 10% formalin for 30 minutes. Immunofluorescence staining was conducted using the SARS-CoV-2 nucleoprotein mouse monoclonal antibody (1:500; Sino Biological #40143-MM05), Alexa-Fluor Plus 488 anti-mouse antibody (1:1,000; Invitrogen #A32723) and DAPI (1:10,000; Invitrogen #D21490). Four images per well were taken at 20x magnification with an EVOS M5000 microscope (Invitrogen, software version 1.3.660.548) for subsequent computational analysis. Image quantification was performed using Cellprofiler 2.2.0 (Carpenter et al., 2006). The analysis pipeline was elaborated and optimised using pictures of cells infected with a blind dilution series of SARS-CoV-2. Rolling-ball background subtraction was performed for both channels using the ImageJ command (Schneider et al., 2012). Given the small cell size, individual cells were segmented based on Otsu thresholding of the DAPI channel. Note that for six wells one picture out of the four pictures taken was excluded from the quantification because of poor nuclei segmentation, as judged by eye. Infected cells were distinguished based on the SARS-CoV-2 N signal (Alexa-Fluor Plus 488 channel) (Supplementary Fig. S4A). To distinguish between infected and uninfected cells, a mean intensity threshold was determined as the 99% quantile of the mean cell intensity in the uninfected control wells of the respective plate (Supplementary Fig. S4B-D). Cells with mean intensity above the threshold were marked as positive and cells below or equal to the threshold were marked as negative (Fig. S4A). The percentage of positive cells per condition was grouped in Python 3.8.4 and plotted against the relative cell density (overall cell count per condition relative to the overall cell count in the infected and DMSO-treated control).

### 2.7. Culture of primary cells

Isolation and establishment of well-differentiated primary human airway epithelial cells was done as described recently (Jonsdottir and Dijkman, 2015). In brief, human tracheobronchial material from explanted human lungs was enzymatically digested for 48h at 4°C prior to isolation of epithelial cells from the lumen. Cells were seeded on collagen-coated transwells with 0.4 µm pore size, polyester membrane inserts (Corning; #3470) and cultured at 37°C and 5% CO2. When confluence was reached, cells were washed with HBSS and exposed to the air. Medium change of the basolateral compartment was performed every other day and cells were washed once weekly with HBSS to remove mucus. Two hours prior to infection, the cells were apically washed with HBSS twice and the basolateral medium was exchanged to medium containing 1 or 5 µM of the respective drug. Lung transplantations were conducted at Hannover Medical School (MHH) in accordance with good clinical and ethical practice. All patients (>18 years of age) gave informed consent for tissue donation and the project was approved by the local ethic committee at MHH (ethic vote 3346/2016). Cells were tested negative for contaminations with mycoplasma (Eurofins).

### 2.8. Pathway analysis

Pathway analysis and acquisition of metadata about the ReFRAME drug repurposing collection was mainly conducted on the basis of information provided by Scripps Research Institute (https://reframedb.org) (Janes et al., 2018) and the Ingenuity Pathway Analysis software suite (IPA, QIAGEN Inc., https://www.qiagenbio-informatics.com/products/ingenuity-pathway-analysis). To complement this information, the following sources were used: DrugBank (https://go.drugbank.com/) (Wishart et al., 2006), PubChem (Kim et al., 2019), Guide to PHARMACOLOGY (Armstrong et al., 2020) and NCBI PubMed (Coordinators, 2018). Data was processed via Microsoft Excel (Version 2016), GraphPad Prism (Version 8) and Adobe Illustrator. All data were acquired in December 2020, data for previous screening results (reframedb.org) were updated in April 2021.

## 3. Results

### 3.1. Profiling of the ReFRAME drug repurposing collection against HCoV 229E, a human-tropic alphacoronavirus

To identify small molecules with broad-spectrum anti-coronavirus activity we used a sequential screening strategy (Fig.1B). First, we screened the comprehensive ReFRAME drug repurposing library with a HCoV 229E *Renilla* luciferase reporter virus (HCoV 229E/R-Luc) using Huh-7.5/F-Luc cells, which are highly permissive for HCoV 229E infection (Supplementary Fig. S1). These cells are engineered to express a firefly luciferase reporter gene, which permits assessment of cell viability and infection efficiency in a dual luciferase reporter assay. In total, 12,993 molecules were analyzed. Using stringent inclusion criteria of ≤6.5% virus infection and >150% cell viability, we identified a total of 137 primary hits (Fig. 2A). We chose a high cell viability threshold, because HCoV 229E/R-Luc infection of these cells is lytic, so that non-toxic, antiviral molecules enhance cell survival. We next validated these candidates using the same cellular system and an 8-step dose series for each molecule to deduce dose-activity relationships for each candidate (Supplementary Fig. S2). This way we confirmed 134 distinct primary hits with measurable IC50 within the dose range tested, corresponding to an overall confirmed hit rate of 1%. According to the ReFRAME database, 115 of these hits were screened against SARS-CoV-2 before (Fig. 2B). Forty-five out of 134 confirmed HCoV 229E hits had emerged as primary hits in a SARS-CoV-2-screening campaign (Riva et al., 2020) and in as yet unpublished studies, making them potential broad-spectrum coronavirus antiviral candidates (Fig. 2B and supplementary table S1). More than 50% of the 134 HCoV 229E hits are either in phase 2 to 3 clinical development, launched, or available as prescription drugs. Metadata of these molecules implicate at least 59 distinct drug target categories, thereby defining a landscape of potential host targets for antiviral therapy against an alphacoronavirus (Fig. 2C, supplementary table S1, and supplementary Fig. S3). Notably, 12 of our 134 confirmed 229E hits were at the same time observed among the 100 confirmed SARS-CoV-2 hits in a screening by Riva et al. making the potential candidate broad-spectrum antivirals against diverse coronaviruses (Fig. 2D) (Riva et al., 2020). The most potent compounds scoring in both screens were Remdesivir, ZLVG CHN2, VBY-825 and Apilimod (Fig. 2E).

**Fig. 2.**
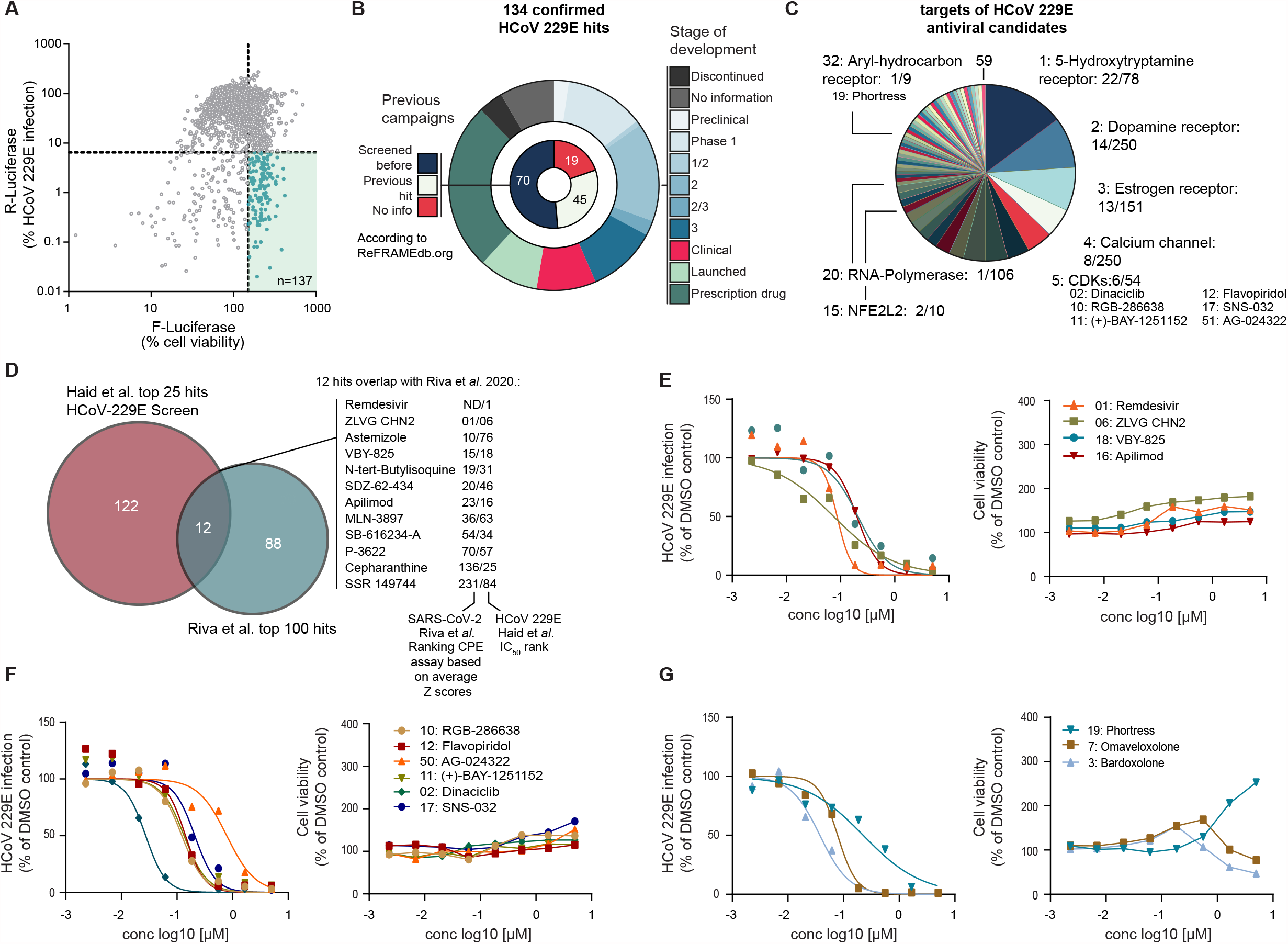
Results of ReFRAME library screening against the alphacoronavirus HCoV 229E. **(A)** Results of the dual luciferase-based HCoV 229E/R-Luc screening of the ReFRAME library in Huh-7.5/F-Luc cells. Renilla luciferase (R-Luc) activity is proportional to virus infection, whereas firefly luciferase (F-Luc) activity corresponds to cell numbers and viability. R-Luc and F-Luc values detected in DMSO-treated HCoV 229E infected cells were set to 100% and corrected for background values in case of R-Luc. **(B)** Stage of development of confirmed HCoV 229E hit compounds and information about previous screens by others. **(C)** Fifty-nine distinct host targets are annotated to these hits. The number before the dash indicates the number of antiviral hits with this target annotation, the number after the dash is equivalent to the total number of molecules within this target category in the ReFRAME library. **(D)** Overlap between this present ReFRAME library screening with HCoV 229E confirmed hits of the previous SARS-CoV-2 screening by Riva at al. (Riva et al., 2020). Numbers indicate the IC50 ranks of the respective studies. **(E)** Dose-dependent antiviral activity of top ranked confirmed hits of Riva et al. against HCoV 229E. Dose-dependent antiviral activity of CDK inhibitors **(F)** and three inflammatory modulators **(G)** against HCoV 229E. Numbers before compound name indicate IC50-rank in the HCoV 229E screening. One experiment with means of two independent luciferase measurements normalized to DMSO control is given.

The most frequent target categories of our confirmed 229E hits were the hydroxytryptamine receptor (22 compounds), the dopamine receptor (14 compounds), and the estrogen receptor (13 compounds) (Fig. 2C). For eight molecules, calcium channels are listed as targets. Notably, six of 54 cyclin-dependent kinase inhibitors (CDKis) included in the ReFRAME library (April 2021) were found to have antiviral activity. Five out of six ranked among the top 50 most active HCoV 229E hits, highlighting the relevance of these kinases for HCoV 229E infection in this cellular system (Fig. 2F). Dinaciclib, a phase III CDKi, exhibited an IC50 of 27.3 nM making it the second most potent compound in the screen, only surpassed by Remdesivir (IC50, 15.7 nM). Notably, Bardoxolone, which is in a clinical trial for treatment of hospitalized COVID-19 patients (BARCONA, ClinicalTrials.gov Identifier: NCT04494646), and Omaveloxolone, both NFE2L2 (also known as NRF2) agonists, ranked at position 3 and 7, respectively (Fig. 2G). Finally, Phortress an aryl-hydrocarbon receptor (AHR) ligand, which also modulates inflammatory processes, inhibited 229E infection with an IC50 of 230nM (Fig. 2G).

### 3.2. Activity of HCoV 229E hits against SARS-CoV-2 infection of Calu-3 cells

As a second step of our screening work flow (Fig. 1B), we determined the potency of these HCoV 229E hits against SARS-CoV-2 (Fig. 3). To this end, we infected the human lung cell line Calu-3 with SARS-CoV-2 (strain SARS-CoV-2/München-1.2/2020/984, p4) (Wolfel et al., 2020) in presence of the 134 HCoV 229E hit substances (screening dose 5 µM). We measured infection by quantification of viral genome equivalents in the culture fluid (Fig. 3A) and by the analysis of total cell and infected cell numbers based on DAPI staining and nucleoprotein immunofluorescence (IF) respectively (Fig. 3B and C). Total cell numbers and frequencies of virus-infected cells were determined in four independent images per well using an automated image processing workflow (Supplementary Fig. S4, and methods). Besides Remdesivir, 12 compounds reduced the virus load in the culture at least two standard deviations below the mean value of the DMSO solvent control (Fig. 3A). Among these candidates, Phortress, an aryl hydrocarbon receptor (AHR) ligand, Flavopiridol, a CDK inhibitor, AXT-914, a calcium-sensing receptor inhibitor, and Gentian violet, an antiseptic dye used for topical treatments of various dermatological conditions, reduced infected cell numbers by more than 50% without reducing total cell numbers to less than 75% (Fig. 3C). Three additional CDKis (Dinaciclib, RGB-286638, (+)-BAY-1251152), and the other candidates, reduced virus load, but did not meet our stringent confirmation criteria based on intracellular N protein staining and cell viability (<50% infected cell, >75% total cells). A survey of these candidates implicated by the qRT-PCR assay and the broad-spectrum antivirals confirmed by the IF-based assay is given in Fig. 3D.

**Fig. 3.**
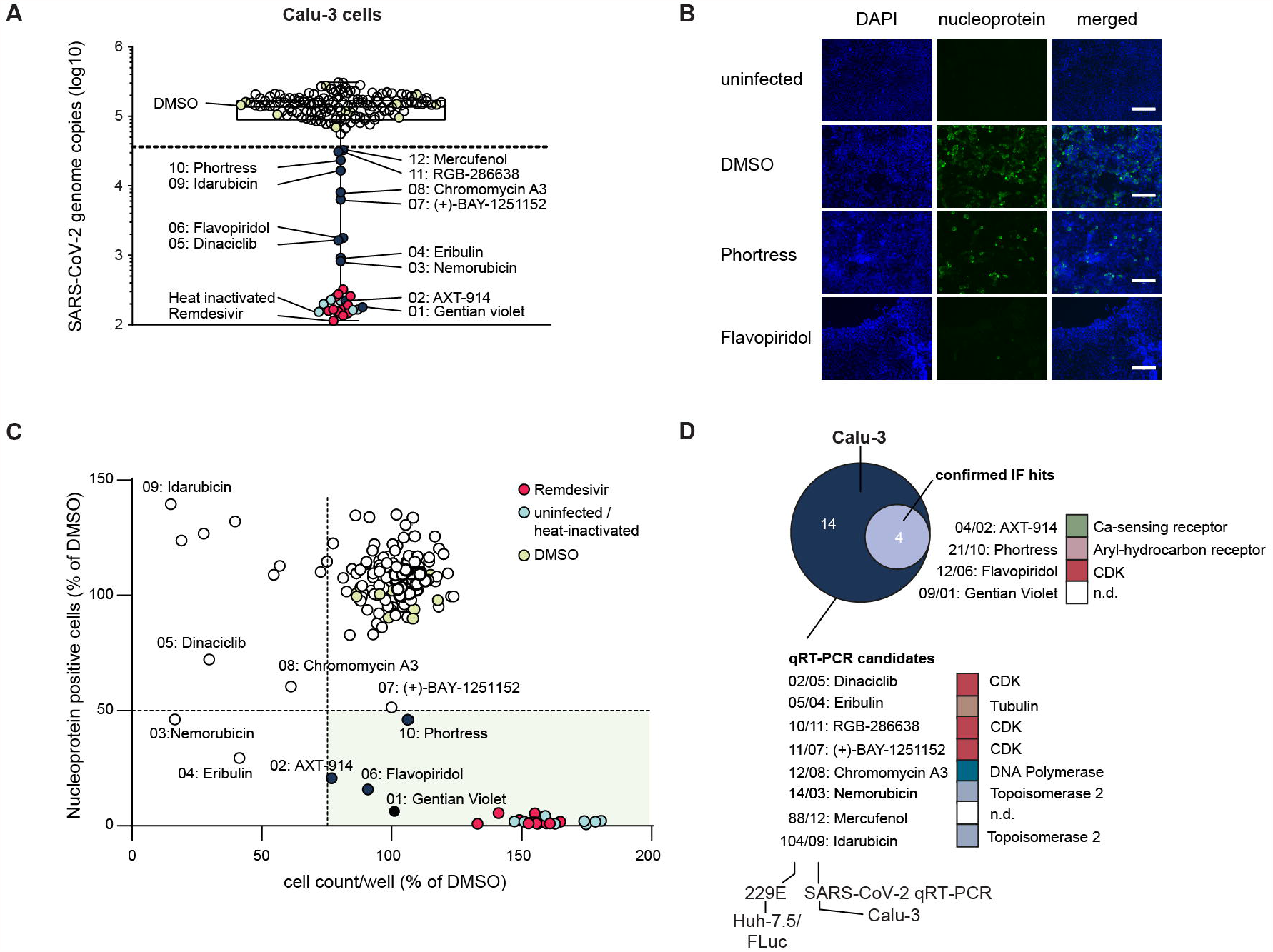
Counter screening of confirmed HCoV 229E hits with SARS-CoV-2. Calu-3 cells were infected with SARS-CoV-2 (strain SARS-CoV-2/München-1.2/2020/984, p4) in the presence or absence of indicated compounds (final concentration 5 µM). **(A)** Infection efficiency was analyzed 48h post inoculation by quantification of viral genome equivalents in the culture fluid of cells. Culture fluid of cells inoculated with heat-inactivated SARS-CoV-2 served as background control (light blue circles). Compounds, which reduced virus load more than two standard deviations below the mean value of the DMSO control, were recognized as candidate antivirals. **(B)** Immunofluorescene analysis of infected cells. Cellular DNA was stained with DAPI (blue), and SARS-CoV-2 nucleoprotein with a monoclonal antibody (green). **(C)** Imaging-based quantification of total cell numbers and infected cell numbers. Images were automatically quantified as outlined in the methods section. The relative number of infected cells (N protein expressing cells) is correlated against the number of total cells (scale bar 125 µm). Means of four images per well of one experiment are given. DMSO and heat inactivated virus controls are depicted with light green and light blue circles, respectively. Data from Remdesivir-treated calls are depicted with red circles. Validated broad-spectrum coronavirus antivirals meeting our inclusion criteria are shown as with dark blue circles. Inclusion criterion was reduction of virus infected cell numbers by more than 50% and residual total cell number of more than 75% of DMSO solvent control. **(D)** Venn diagram highlighting the total number of qRT-PCR-based broad-spectrum candidates and confirmed hits based on the immunofluorescence analysis. Annotated targets of molecules with broad activity against both HCoV 229E and SARS-CoV-2 are given at the right. n.d., no data.

### 3.3. Confirmation of broad-spectrum anti-coronavirus drug candidates in well differentiated human airway epithelial air liquid interface cultures

We obtained commercially available candidates with HCoV 229E and SARS-CoV-2 antiviral activity in at least one of the human cell lines and tested their antiviral effect on SARS-CoV-2 infection in well-differentiated primary human airway epithelial cells from three independent donors (Fig. 4). Viral genome copy numbers in cell extracts (Fig. 4A) were consistent with viral copy numbers in the culture fluid (Fig. 4B) and confirmed that Remdesivir and further 10 of 12 tested compounds dose-dependently reduced virus load. However, concomitantly, GAPDH mRNA expression was reduced in case of CDK inhibitors BAY-1251152, Flavopiridol, RGB-286638, and Dinaciclib, suggesting that compromised cell viability may at least partially account for attenuation of virus propagation (Fig. 4C). Likewise, treatment with Nemorubicin, a topisomerase 2 inhibitor, reduced virus load but also GAPDH mRNA expression. Gentian violet and Idarubicine did not reduce virus infection of primary human lung cells. In contrast, Bardoxolone, Omaveloxolone and Phortress displayed potent antiviral effects and modest effects on GAPDH mRNA at a dose of 1 µM. AXT-914, which inhibited both HCoV 229E and SARS-CoV-2 was not commercially available for confirmatory assays in primary lung cells.

**Fig 4.**
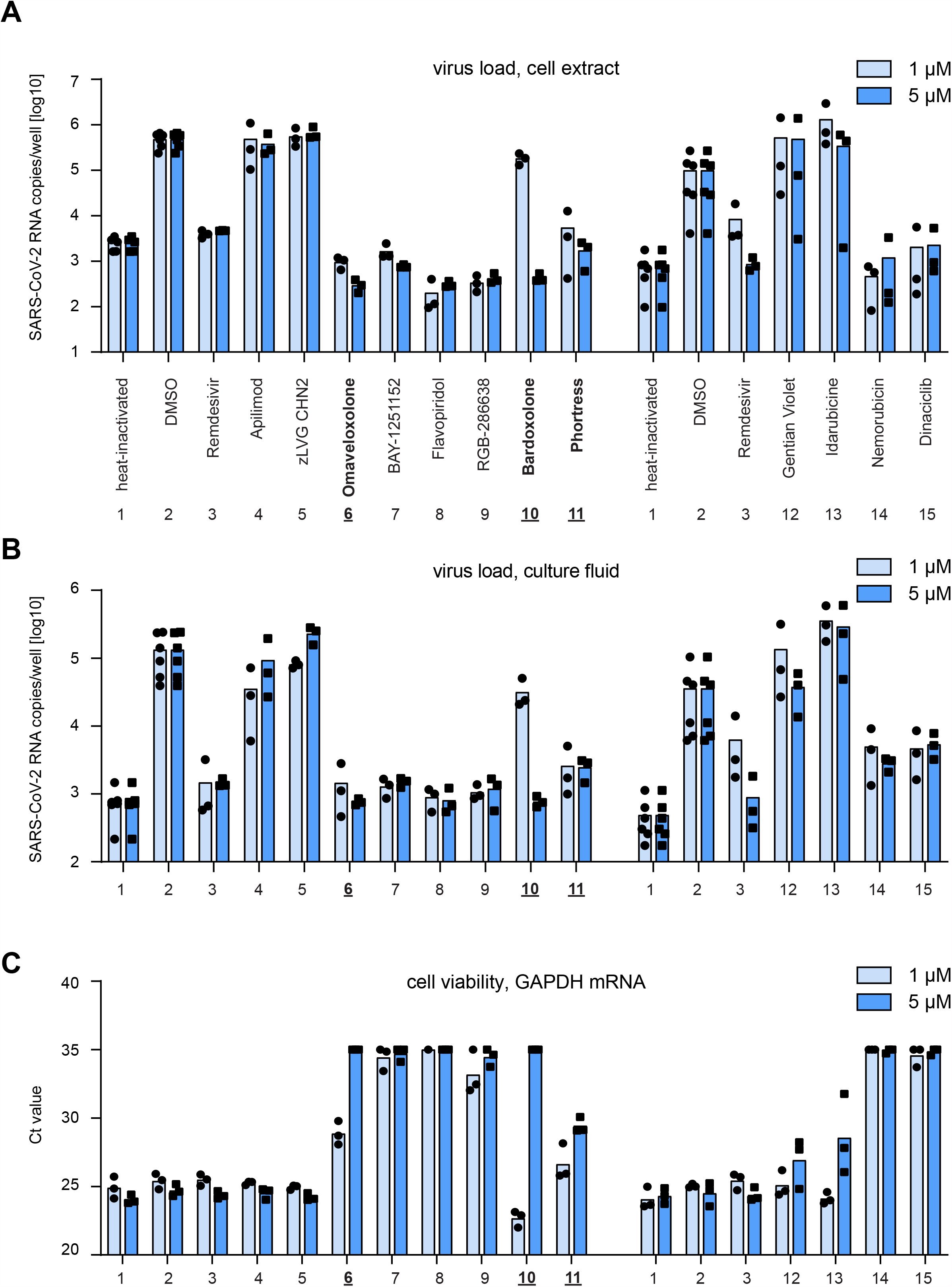
Validation of candidates with broad-spectrum coronavirus activity in well differentiated primary human lung cells. Primary cells from three independent donors were differentiated and grown as pseudostratified epithelium in air liquid interface culture. Cells were pre-treated for 2h with final concentrations of 1 µM (bright blue) or 5 µM (dark blue) of compounds from the basolateral side prior to inoculation with SARS-CoV-2 in the presence of compounds (apical and basal treatment). Infection efficiency was determined 72h post inoculation by qRT-PCR of cell extracts **(A)** and culture fluids **(B). (C)** Cell viability was assessed by quantification of GAPDH mRNA. Ct values >35 were excluded from the analysis. Mean values (bar) and individual values (points) of three donors are given. Analysis was performed in two independent experiments (as indicated by the separation). In both experiments, primary cells from three different donors were used. Virus stocks differed in titer between these experiments (see Material and Methods).

## 4. Discussion

This study provides a comprehensive survey of molecules from the ReFRAME repurposing library with antiviral activity against a human pathogenic alphacoronavirus (HCoV 229E). Combined with a counter-screening using a recent SARS-CoV-2 isolate representing the genus *Betacoronavirus* we highlight compounds with potentially broad-spectrum activity against diverse coronaviruses. Based on access to ample metadata for many of these molecules and the fact that this library accumulates compounds modulating physiological processes in humans, this provides a unique overview of potentially druggable host pathways and targets for development of broad-spectrum coronavirus antivirals.

Primary screening of 12,993 library compounds and dose-response titrations identified and confirmed 134 candidates against HCoV 229E implicating 59 different potential host target categories. Molecules with the 5-hydroxytryptamine (i.e. serotonin) receptor as target annotation represented the largest group of hits against HCoV 229E. We identified 22 of 78 molecules included in the ReFRAME library representing 28% of compounds of this class. The second most dominant HCoV 229E target category was the dopamine receptor with 14 hits out of 250 molecules contained in the library. Thus, collectively, 36 of 134 HCoV 229E targeting compounds, equivalent to 26.8%, are categorized as neurotransmitter receptor modulators (target information: reframedb.org, February 2021). Importantly, only one molecule (SB-616234-A) has been described in a previous ReFRAME SARS-CoV-2 screening study (Riva et al., 2020), and none of them scored in our SARS-CoV-2 counter screening, suggesting that these compounds and their targets are specific for HCoV 229E. The 5-hydroxytryptamine receptor family comprises seven protein groups, six of which encompass G-protein coupled receptors (GPCRs) whereas the dopamine receptor family includes proteins categorized in two groups of GPCRs (Pytliak et al., 2011). Therefore, GPCR-targeting compounds are heavily enriched in the HCoV 229E antiviral screening. The reason for the HCoV 229E preferential accumulation of GPCR modulators is currently unknown but likely due to key differences in host factor usage between these viruses.

One important difference between HCoV 229E and SARS-CoV-2 is their receptor usage with the former exploiting human alanyl aminopeptidase (ANPEP, also known as hAPN/CD13) and the latter ACE2. Both proteins are plasma membrane resident proteases, but neither is a GPCR. ANPEP is expressed in numerous tissues and hydrolyses proteins involved in different physiological processes (Luan and Xu, 2007). Intestinal ANPEP processes peptides originating from gastric proteases, whereas neuronal ANPEP for instance is involved in the proteolysis of neurotransmitters. It will be interesting to explore if compounds from the 5-hydroxytryptamine receptor and dopamine receptor categories inhibit ANPEP-dependent cell entry, and if so, by which mechanism this occurs. GPCR-targeting neuroleptic drugs have emerged as cell entry inhibitors of other enveloped viruses before, although these viruses also do not use GPCRs directly for cell entry either (Pietschmann, 2017). The mechanisms by which this occurs may differ between these viruses and may range from direct binding to the viral fusion protein to the modulation of membrane curvature and fluidity.

Next to Remdesivir, the CDKi Dinaciclib emerged as the most potent confirmed HCoV 229E inhibitor within the ReFRAME library (IC50 27 nM). We identified five additional CDKis, all of which ranked among the top 50 HCoV 229E screening hits with an antiviral activity well-separated from cytotoxicity (Fig. 2F and supplementary Fig. S2). Thus, the CDKi category hosts the most potent HCoV 229E inhibitors in the Huh-7.5/F-Luc assay. CDKis were not described as SARS-CoV-2 inhibitors by Riva et al. (Riva et al., 2020), possibly due to usage of different host cell lines. However, a recent phosphoproteome survey of SARS-CoV-2 infected cells has revealed a time-structured, SARS-CoV-2-dependent modulation of CDK activity (Bouhaddou et al., 2020). These authors also described several potential CDK phosphorylation sites within viral proteins and they found that Dinaciclib inhibits SARS-CoV-2 infection of Vero E6 and of A549/ACE2 cells with nanomolar efficacy (Bouhaddou et al., 2020). Flavopiridol, a phase 3 CDKi, qualified as a hit in both imaging-and qRT-PCR-based assays in Calu-3 cells. RGB-286638, (+)-BAY-1251152, and Dinaciclib also reduced SARS-CoV-2 RNA copy numbers in the culture fluid of infected Calu-3 cells, but they did not meet the more stringent inclusion criteria of the Calu-3 imaging-based analysis (Fig. 3). CDKs are essential regulators of cell cycle progression so that unwanted side effects need careful consideration. In the 8-step dose-activity titration of these compounds in Huh-7.5/F-Luc cells, we observed a clear separation between antiviral activity against HCoV 229E and cytotoxicity, suggesting a direct inhibition of virus infection. Using a limited dose range in the human primary lung cells, we were unable to separate antiviral effects from cytotoxic effects. Thus, CDK inhibition seems broadly active against coronaviruses in some cell lines and inhibition of these kinases merits further attention for development of broad-spectrum coronavirus inhibitors. Further to this, Phortress, an AHR ligand, emerged as broad-spectrum antiviral with activity in primary differentiated human lung cells. Ligands of the AHR are in clinical development for treatment of kidney cancer (Itkin et al., 2020) (ClinicalTrials.gov Identifier: NCT04069026).AHR itself is a transcription factor that resides in the cytoplasm complexed with the heatshock protein 90 (Hsp90). Upon ligand binding, AHR disassociates from Hsp90, traffics to the nucleus and transactivates genes downstream of the xenobiotic response element. AHR-dependent genes include metabolic enzymes such as cytochrome (CYP) P450 1A1, CYP1A2 and CYP1B1 (Itkin et al., 2020). AHR also participates in immune regulatory processes: for instance constitutive activation of AHR downregulates type I IFN-dependent innate immune responses (Yamada et al., 2016). Notably, Kueck *et al*. reported that AHR activation in macrophages limits cellular dNTP pools, thereby inhibiting replication of herpes-and retroviruses (HSV-1, and HIV-1, respectively) (Kueck et al., 2018).

Finally, Bardoxolone and Omaveloxolone exhibited an IC50 of 37 nM and 78 nM against HCoV 229E, ranking them at position 3 and 7, respectively, in our hit list. While they did not meet inclusion criteria of our SARS-CoV-2 Calu-3 cell counter screen, they exhibited an antiviral activity against SARS-CoV-2 in primary human lung cells. Both compounds are semisynthetic derivatives of the triterpene oleanolic acid. They bind covalently to Kelch-like ECH-associated protein 1 (KEAP1), which leads to the activation of NFE2L2 (Nrf-2). Because of its anti-inflammatory and tissue repair effects, NFE2L2 activation has been recently proposed as a strategy to treat COVID-19 (Cuadrado et al., 2020) and a clinical trial testing safety, tolerability, and efficacy of Bardoxolone methyl in hospitalized COVID-19 patients is ongoing (BARCONA, ClinicalTrials.gov Identifier: NCT04494646). In fact, broad-spectrum antiviral properties of Bardoxolone methyl have been reported for a variety of viruses including dengue virus (DENV), Zika virus (ZIKV), hepatitis B virus (HBV), hepatitis C virus (HCV) and herpes simplex virus 1 (HSV-1) (Beigel et al., 2020; Nio et al., 2019; Rothan et al., 2019). Specifically, the NFE2L2-dependent expression of antioxidant genes was found to be suppressed in biopsy samples obtained from COVID-19 patients, and the administration of the primary metabolite prodrugs 4-octyl-itaconate (4-OI) and dimethyl fumarate (DMF), that both activate NFE2L2, had anti-SARS-CoV-2 effects at high concentrations (Olagnier et al., 2020). Bardoxolone methyl has progressed to phase 3 clinical trials, mainly as a long-term treatment of kidney diseases. Although its late stage clinical development faced setbacks due to the occurrence of cardiovascular side effects in the BEACON study (de Zeeuw et al., 2013), the mechanisms and safety biomarkers for such side effects have been identified, and the clinical development of Bardoxolone methyl has been resumed (Kanda and Yamawaki, 2020). In addition, Omaveloxolone is in late-stage clinical trials for a neurological indication (Lynch et al., 2021). Due to the existence of ample clinical safety data even as longer term treatment, a mechanistic rationale, and the prospect to exert both antiviral and host-protective effects, the repurposing of NFE2L2 activators as broad-spectrum anti-corona drugs merits particular attention.

## 5. Conclusions

This two-step screening of the ReFRAME repurposing library provides a comprehensive overview of drug-(like) molecules with antiviral activity against the human pathogenic alphacoronavirus HCoV 229E. It implicates 59 distinct cellular targets as potential host targets for inhibition of HCoV 229E infection. The counter-screening of 134 confirmed HCoV 229E antivirals revealed four compounds with antiviral activity against both HCVo 229E and SARS-CoV2 infection of Calu-3 cells (AXT-914, Phortress, Flavopiridol, Gentian violet). Next to Flavopiridol, additional CDKi inhibited SARS-CoV2 infection albeit with less favorable cytotoxicity profile, suggesting that side effects of these molecules will limit utility as coronavirus antivirals. Phortress, Bardoxolone and Omaveloxolone, emerged as candidate broad-spectrum coronavirus antivirals with moderate safety profile in primary human lung cells. Thus, modulation of AHR-and/or NFE2L2-dependent host responses merits attention for development of broad-spectrum antivirals against coronaviruses.

## Acknowledgments

We are grateful for the gift of the SARS-CoV-2/München-1.2/2020/984 strain by Christian Drosten and to Volker Thiel for sharing the HCoV 229E/R-Luc reporter virus. This publication was supported by the European Virus Archive GLOBAL (EVA-GLOBAL) project that has received funding from the European Union’s Horizon 2020 research and innovation programme under grant agreement No. 871029. We thank Charles M. Rice for the gift of Huh-7.5 cells, Gert Zimmer for 16HBE140 and Volker Thiel for MRC-5 cells. We also gratefully acknowledge gifts of Vero and Calu-3 cells by Stefan Pöhlmann, and provision of the TCID50-calculator by Marco Binder, and Anne Kühnel and Xiaoyu Zhang for technical support.

## Appendix A. Supplementary data

Supplementary data to this article can be found online.

## Authors’ contributions

Conceptualization: S.H., T.P.

Methodology: S.H., A.M., M.W., A.P.G., S.M.S., L.L., F.Z., G.B., G.G.

Validation: S.H., A.M., M.W., J.R., A.P.G.

Formal analysis: S.H., A.M., M.W., S.M.S., A.P.G., D.K., G.V.

Investigation: S.H., A.M., M.W., J.R., A.P.G., L.L., F.Z

Resources: B.W., T.T.N., U.B., T.F.S.

Data curation: S.H., A.M., M.W., S.M.S., A.P.G.

Writing – original draft: S.H., A.M., M.W., A.P.G., G.V., M.B., T.P.

Writing – review & editing: S.H., A.M., M.W., S.M.S., M.B., A.P.G., L.L., D.K., G.V., B.W., F.Z., G.B., G.G., U.B.

Visualization: S.H., A.M., M.W., S.M.S., D.K., T.P.

Supervision: G.V., G.G., T.F.S, T.P. Project administration: S.H., M.B., T.P.

Funding acquisition: M.B., T.F.S., T.P., G.V., G.G., U.B.

