## Supplementary figures and images for "Repurposing screen highlights broad-spectrum coronavirus antivirals and their host targets"

### Supplementary Figure 1

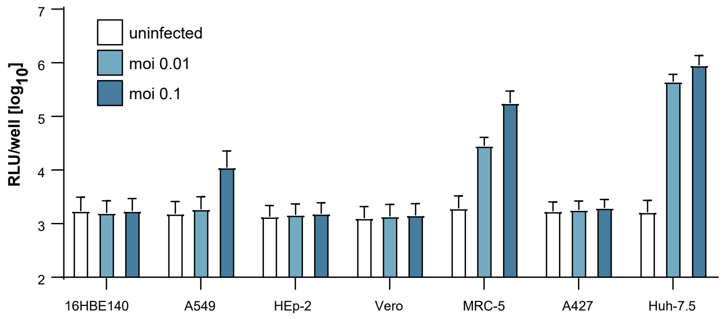

### Supplementary Figure 2

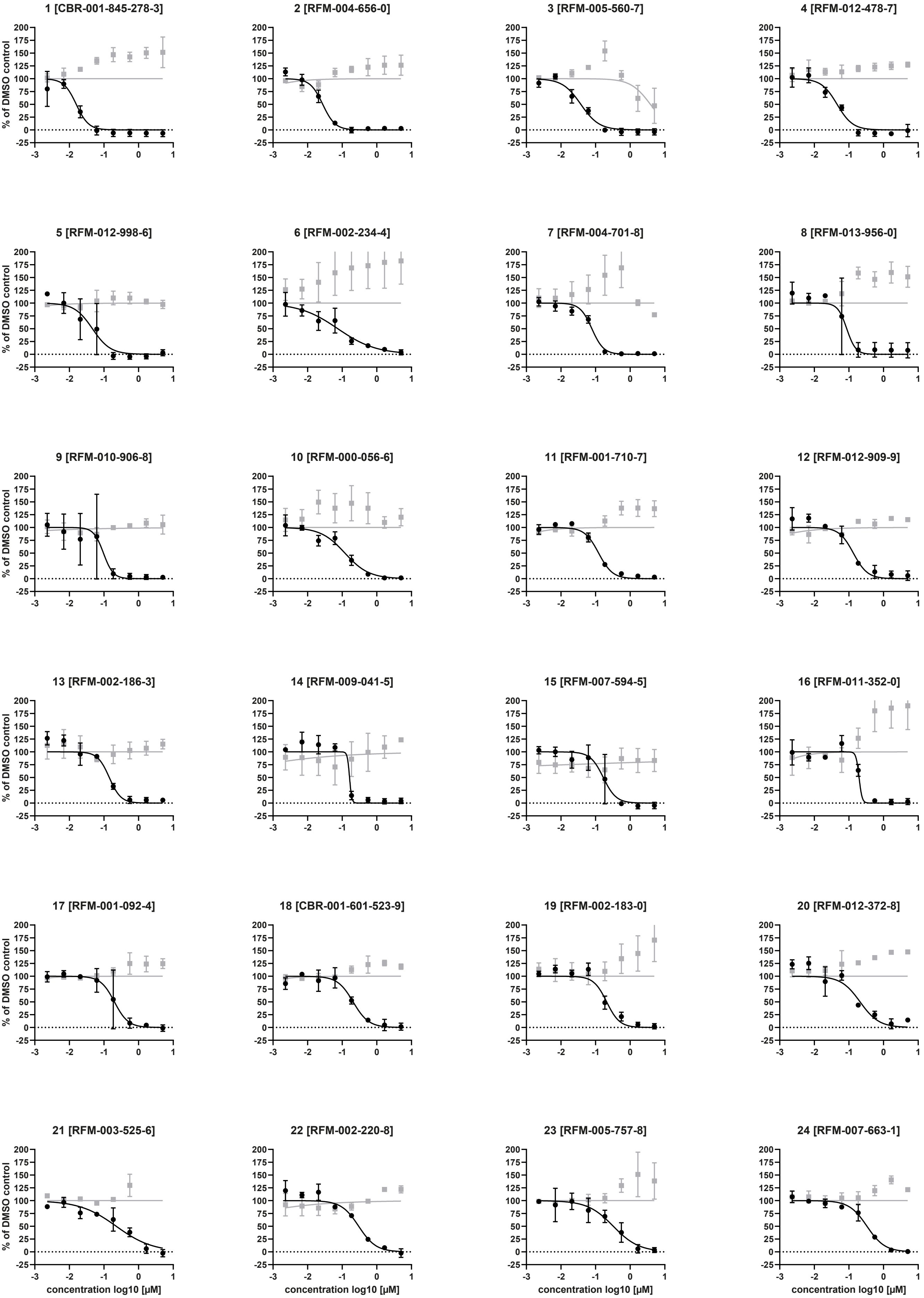

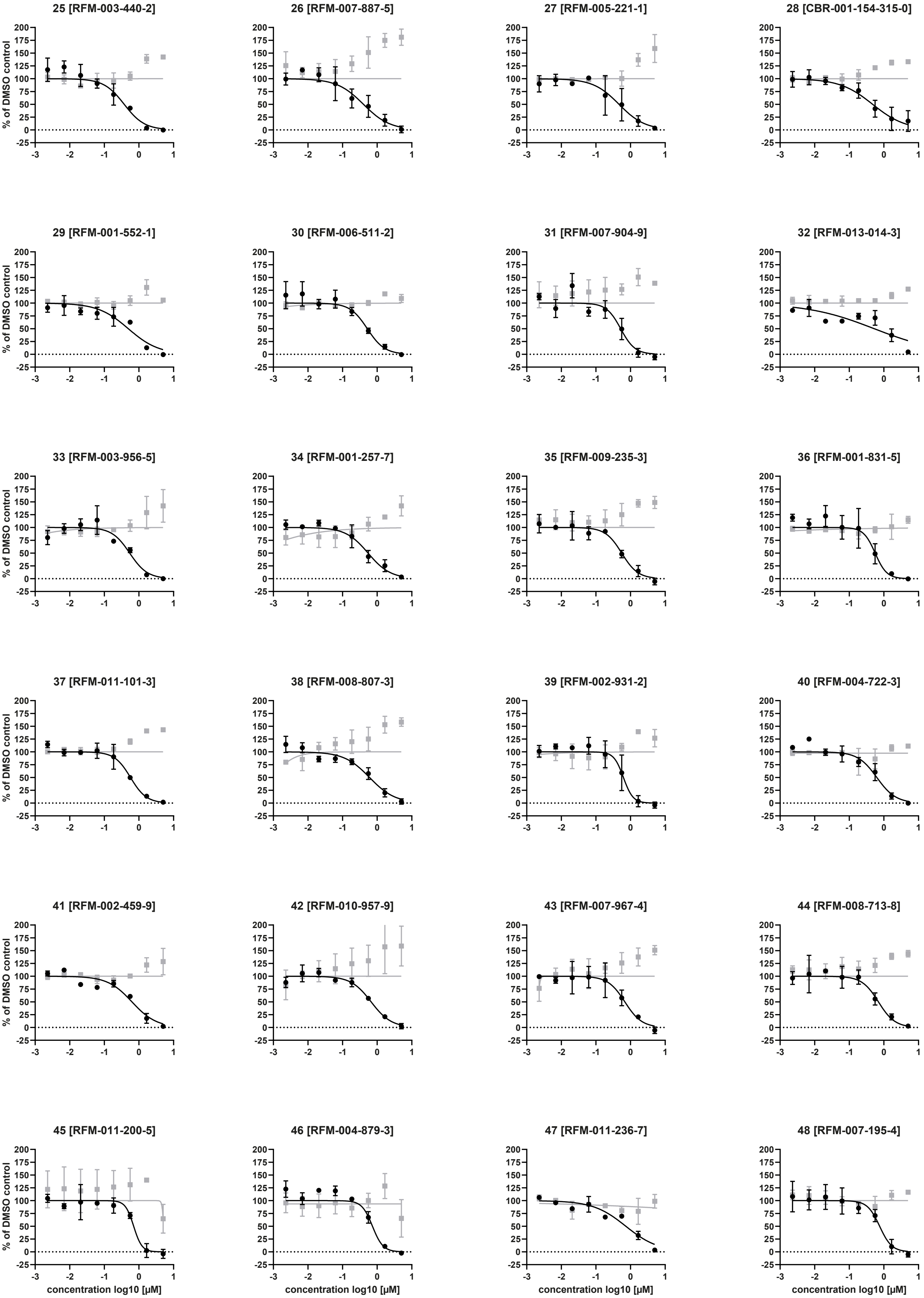

49 [RFM-010-140-6]

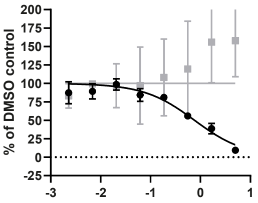


50 [RFM-007-890-0]

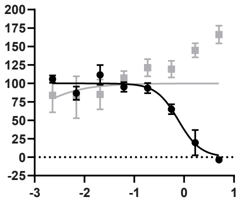


51 [RFM-001-563-4]

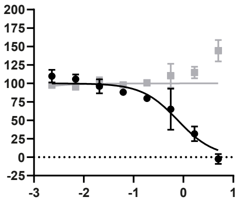


52 [RFM-011-405-6]

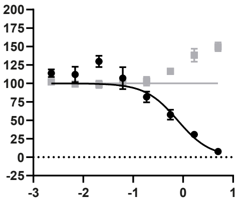


53 [RFM-011-216-3]

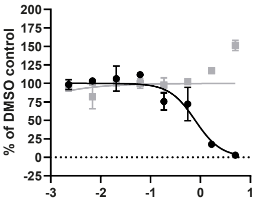


54 [RFM-001-557-6]

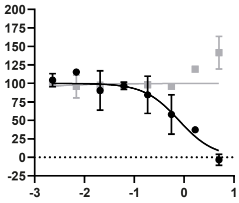


55 [RFM-011-199-9]

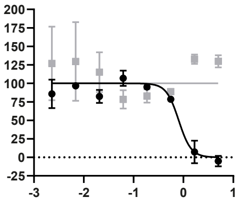


56 [RFM-008-473-1]

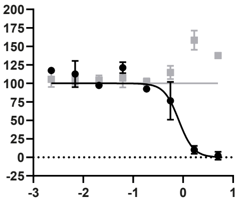


57 [RFM-008-636-2]

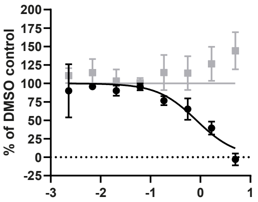


58 [RFM-010-543-1]

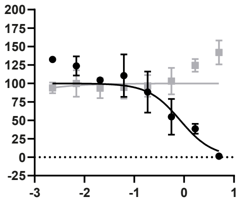


59 [RFM-011-108-0]

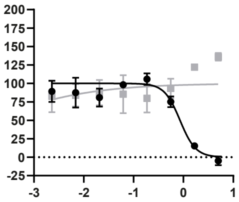


60 [RFM-009-175-8]

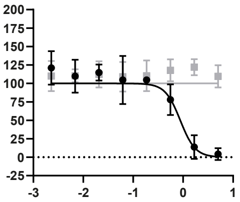


61 [RFM-011-474-9]

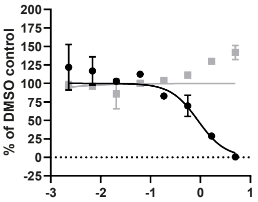


62 [RFM-002-372-3]

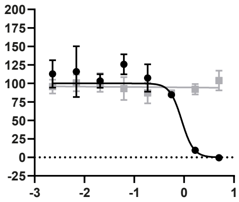


63 [RFM-010-777-7]

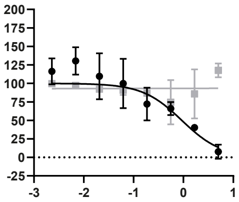


64 [RFM-000-628-0]

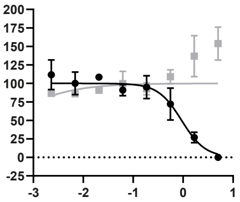


65 [RFM-004-358-3]

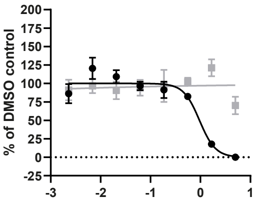


66 [RFM-010-993-3]

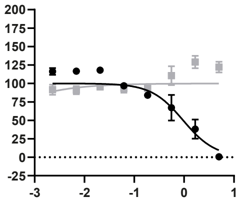


67 [RFM-000-077-1]

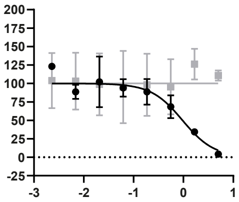


68 [RFM-006-751-6]

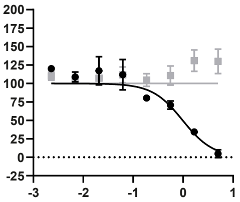


69 [RFM-008-367-0]

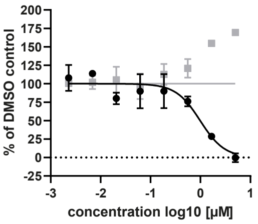


70 [RFM-001-499-3]

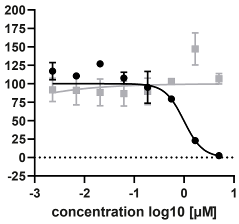


71 [RFM-004-033-5]

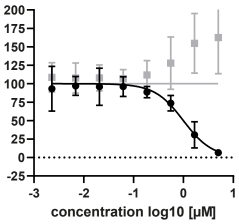


72 [CBR-001-210-396-7]

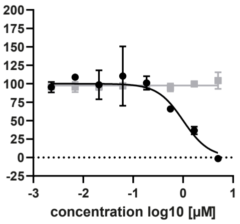

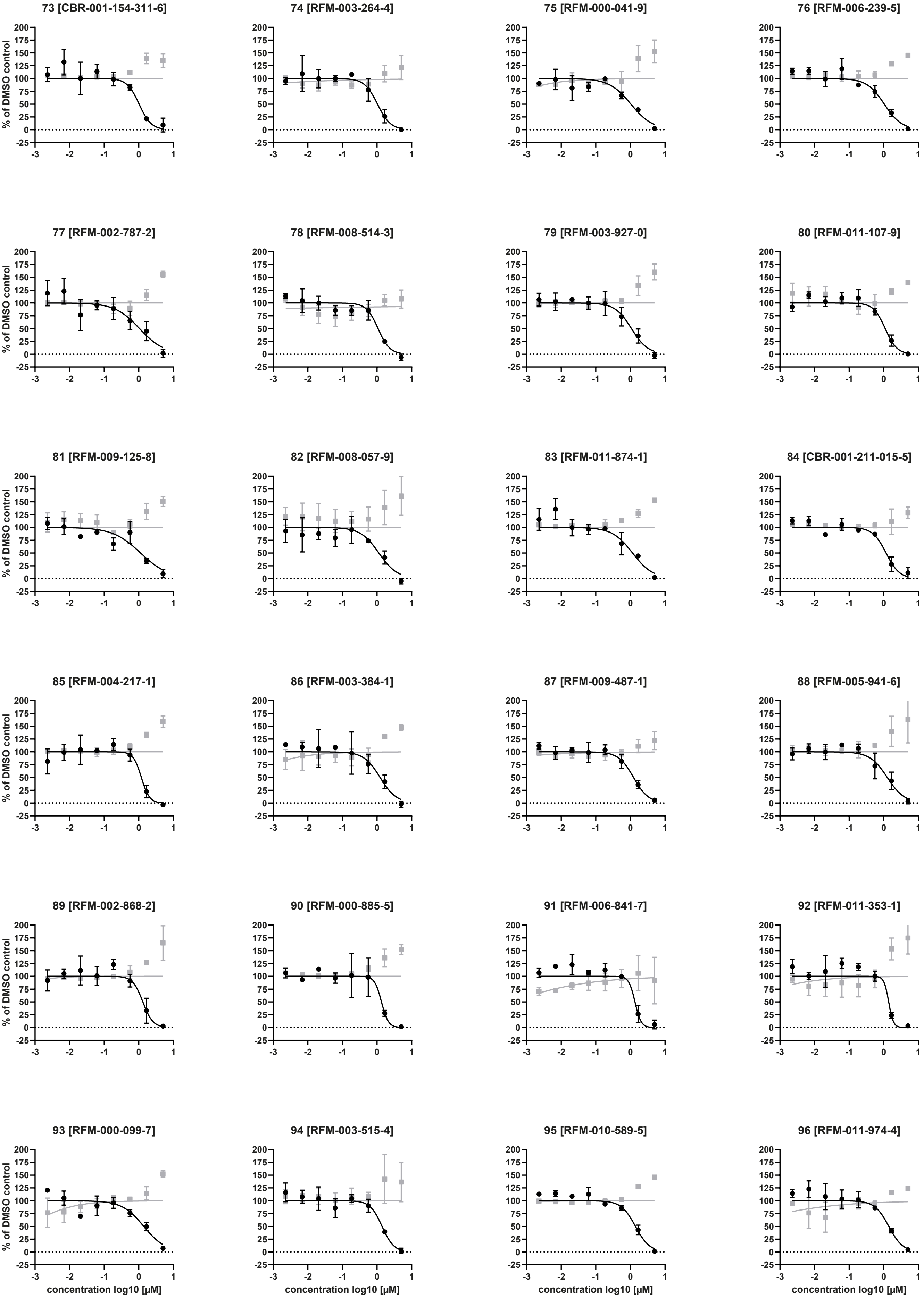

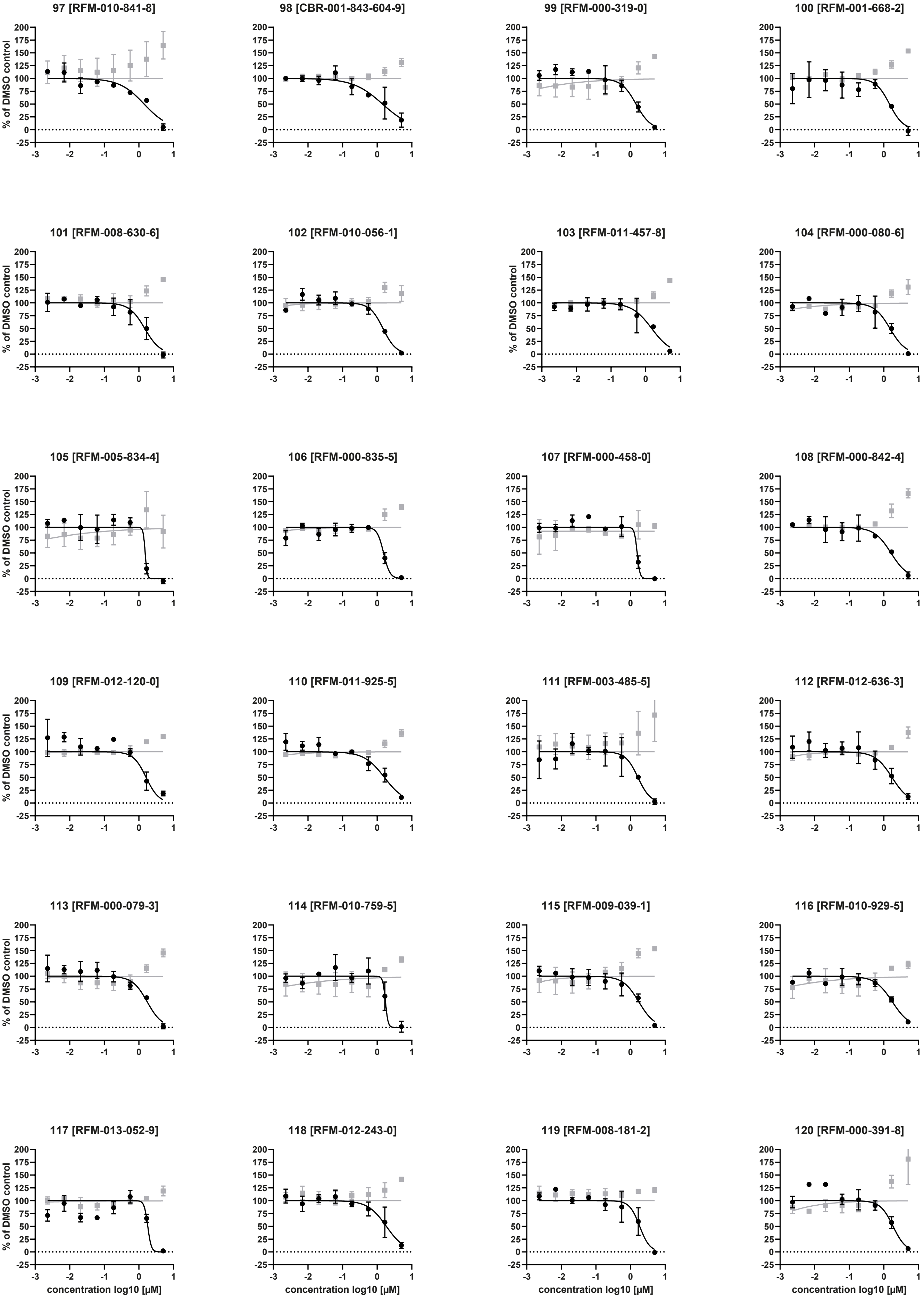

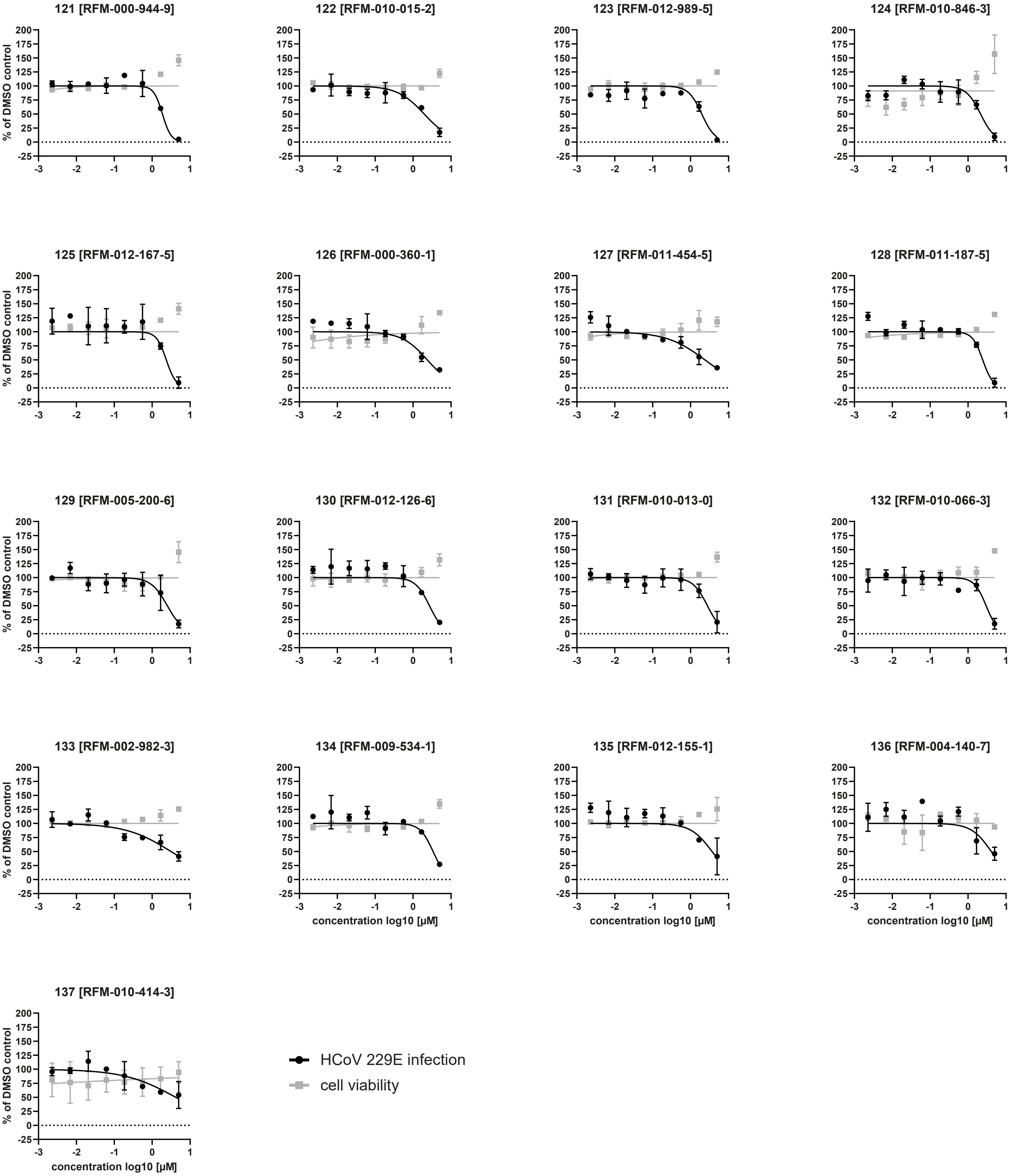

### Supplementary Figure 4

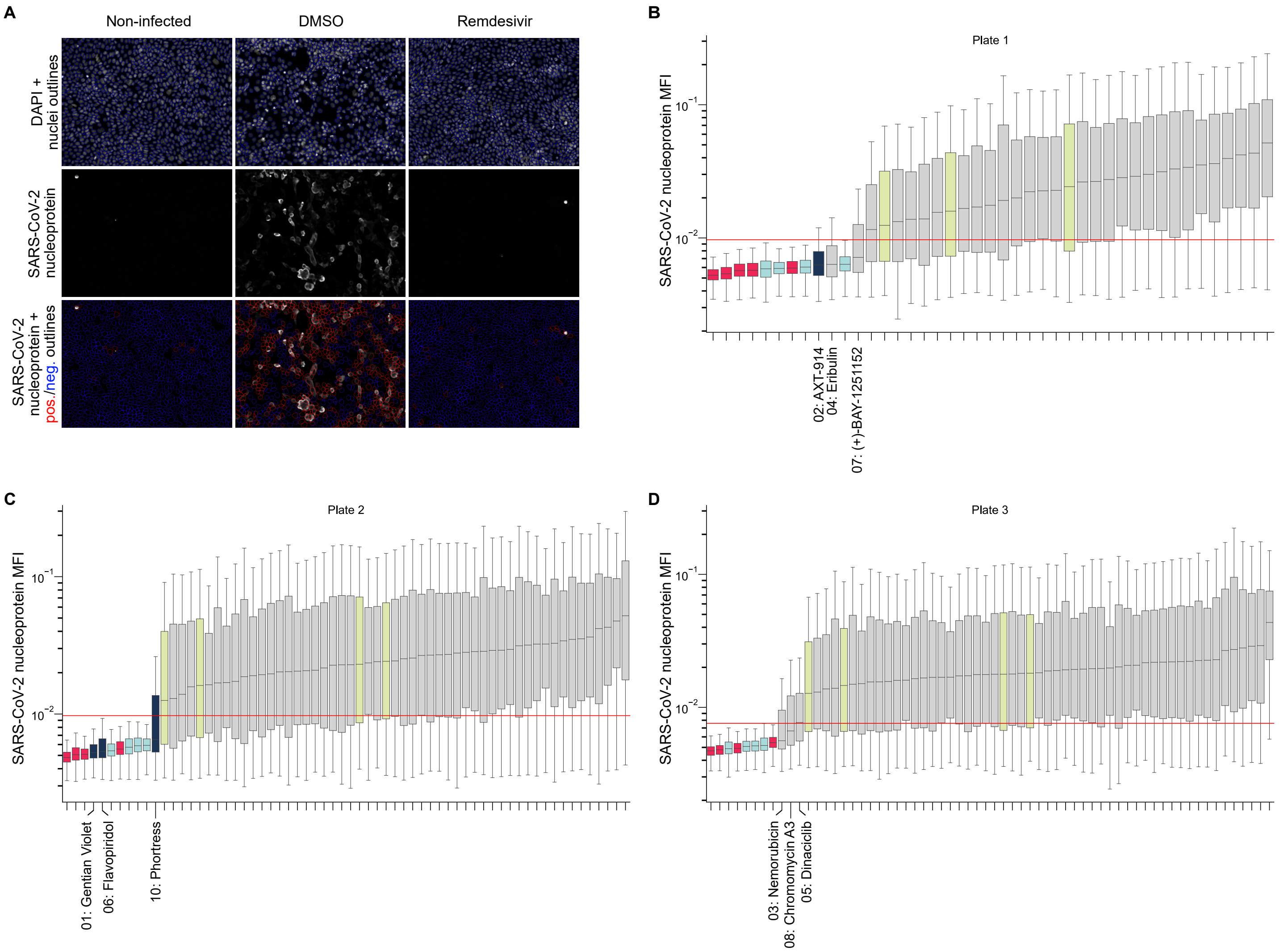
