## Supplementary Figure 3 for "Repurposing screen highlights broad-spectrum coronavirus antivirals and their host targets"

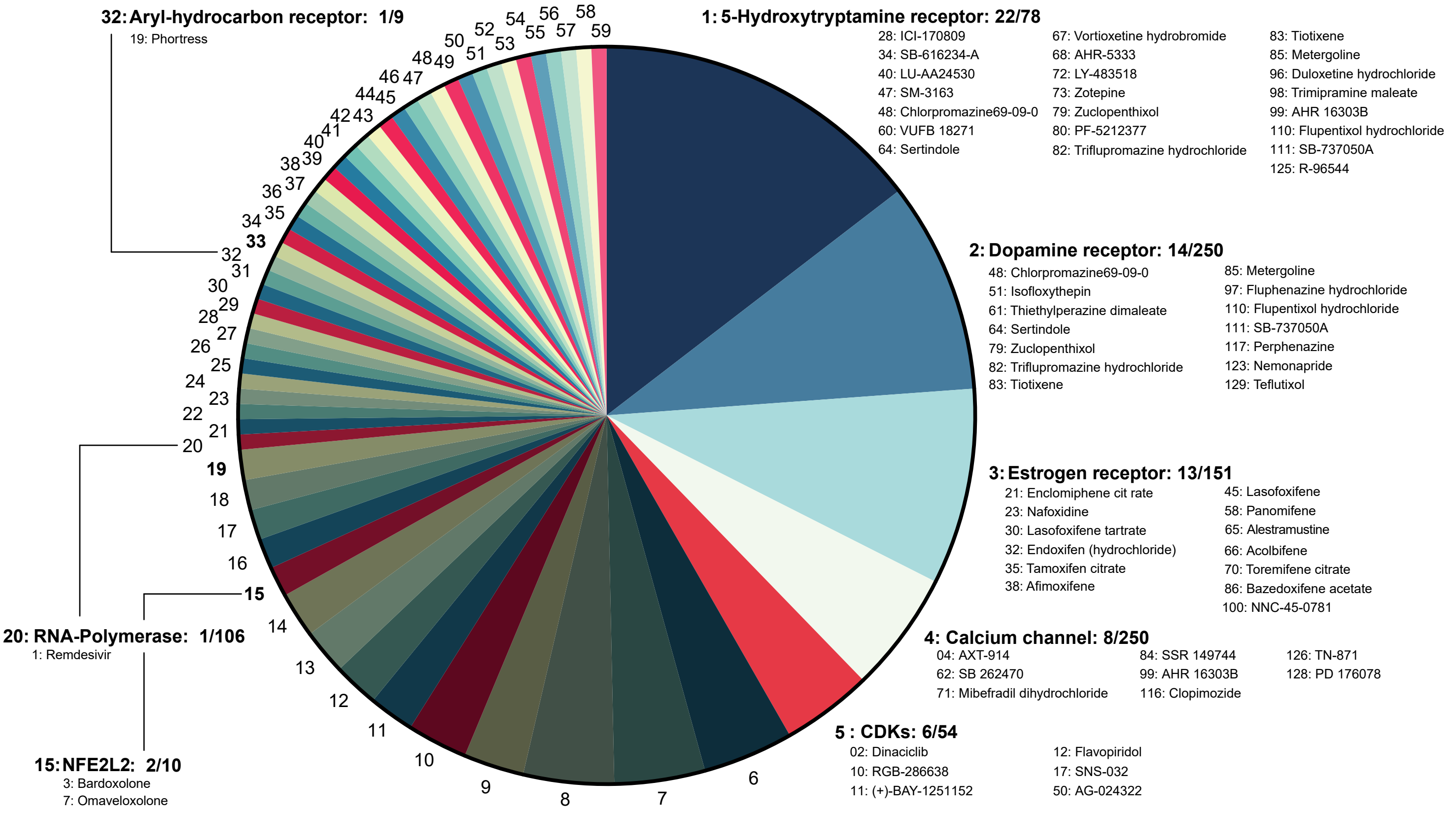


1 5-hydroxytryptamine receptor
2 Dopamine receptor
3 Estrogen receptor
4 Calcium channel
5 Cyclin dependent kinases CDKs
6 Adrenergic receptor
7 Solute carrier family
8 Histamine receptor
9 Tubulin
10 Potassium channel
11 Tachikinin receptor
12 Topoisomerase (DNA)
13 CYP
14 Protein kinase C (PKC)
15 NFE2L2
16 ABCB
17 Squalene synthetase
18 Calmodulin
19 NF-KB
20 RNA polymerase
21 PPARG
22 LONP
23 CaSR
24 Rhinovirus 3C protease
25 Cystein Proteinase
26 Hif
27 Guanine-cytosine (G-C) base pairs
28 Cholecystokinin receptor
29 Interleukin/interleukin receptor
30 PIKFYVE
31 Cathepsin
32 Aryl-hydrocarbon receptor
33 Phospholipase
34 Androgen receptor
35 Prolyl endopeptidase
36 Importin
37 Heat shock protein
38 ABCC
39 Thrombin
40 Thrombin receptor
41 NMDA receptor
42 Bradikinin receptor
43 PIM
44 Steroid receptor
45 Smoothened (SMO)
46 Carnitine palmitoyltransferase (CPT)
47 Serine/threonine kinase AKT
48 Platelet-activating factor receptor (PAFR)
49 Calcium sensing receptor
50 microtubule-associated proteins (MAPs)
51 Sigma receptor
52 Lipogygenase
53 Cyclooxygenase
54 Sphingosine-phosphate receptor
55 Muscarinic acetylcholine receptor (CHRM)
56 Myosin light chain kinase (skeletal/cardiac muscle)
57 μ-Opioid receptor
58 Tyrosin-protein kinase SYK
59 Vasopressin (AVP) V1b receptor
